# Carboxylic Acids that Drive Mosquito Attraction to Humans Activate Ionotropic Receptors

**DOI:** 10.1101/2022.10.20.513041

**Authors:** Garrett Ray, Robert M. Huff, John S. Castillo, Anthony J. Bellantuono, Matthew DeGennaro, R. Jason Pitts

## Abstract

The mosquito, *Aedes aegypti*, is highly anthropophilic and continues to transmit debilitating arboviruses within human populations. Female mosquitoes are attracted to sources of blood by responding to odor plumes that are emitted by their preferred hosts. Acidic volatile compounds, including carboxylic acids, represent particularly salient odors driving this attraction. Importantly, carboxylic acids are major constituents of human sweat and volatiles generated by skin microbes. As such, they are likely to impact human host preference, which is a dominant factor in disease transmission cycles. A more complete understanding of mosquito host attraction will necessitate the elucidation of molecular mechanisms of volatile odor detection that function in peripheral sensory neurons. Recent studies have shown that members of the variant ionotropic glutamate receptor gene family are necessary for physiological and behavioral responses to acidic volatiles in *Aedes*. In this study, we have identified a subfamily of variant ionotropic receptors that share sequence homology across several important vector species and are likely to be activated by carboxylic acids. Moreover, we demonstrate that selected members of this subfamily are activated by short chain carboxylic acids in a heterologous cell expression system. Our results are consistent with the hypothesis that members of this receptor class underlie acidic volatile sensitivity in vector mosquitoes and provide a frame of reference for future development of novel mosquito attractant and repellent technologies.

## Introduction

Mosquitoes, like other insects, possess the remarkable innate ability to sense and respond to multimodal environmental stimuli. Female mosquitoes utilize a variety of transmembrane sensory receptors and specialized innervated sensory structures to detect sensory inputs such as carbon dioxide (CO_2_), airborne odors, heat, moisture and visual cues for the purpose of resource acquisition (Sorrells *et al*. 2022; Raji & DeGennaro 2017; Sparks & Dickens 2016; Ray *et al*. 2014; Suh *et al*. 2014). Volatile organic compounds (VOCs) serve as important kairomones for female mosquitoes, signaling the presence and identities of potential bloodmeal hosts (McBride 2016; Zwiebel & Takken 2004; Takken & Knols 1999). Because mosquito-borne arbovirus transmission depends on female blood feeding on susceptible, permissive hosts, the molecular processes underlying this complex behavior are of keen interest (Huang *et al*. 2019; Schneider & Higgs 2008; Day 2005). This is especially relevant in species such as *Aedes aegypti*, one of the major vectors of Dengue, Zika, and Yellow Fever, that display a high degree of anthropophily (McBride 2016).

Large families of chemoreceptors that are encoded in the genomes of mosquitoes comprise the odorant receptors (ORs), gustatory receptors (GRs), and ionotropic receptors (IRs) (Carey *et al*. 2010; Benton *et al*. 2009; Kent *et al*. 2008). Chemoreceptors are expressed in sensory neurons (Lombardo *et al*. 2017; Matthews *et al*. 2016) and, together with concomitant anatomical structures in the antennae, maxillary palps, labella, and tarsi, provide the molecular and physiological basis for chemical sensitivities (Suh *et al*., 2014; Kaupp, 2010). Characterizing the function of chemoreceptors by identifying ligand-receptor pairs is an important prelude to genetic, neurophysiological, and behavioral studies that will address the systematic effects of both unitary chemical compounds and odor blends on mosquito host choice.

Carboxylic acids (CAs) are a major constituent of human sweat and microbial-produced skin emanations and are known to increase anthropophilic vector species’ attraction to humans (De Obaldia *et al*. 2022; Lucas-Barbosa *et al*. 2022; Showering *et al*. 2022; Ellwanger *et al*. 2021; Smallegange *et al*. 2011; Bernier *et al*. 2007; Smallegange *et al*. 2005; Braks *et al*. 2001; Cork & Park 1996). Previous molecular, electrophysiological, and behavioral studies have demonstrated that members of the IR family of chemoreceptors are involved in the detection of acidic volatiles in mosquitoes (Pitts *et al*. 2017; Raji *et al*. 2019). A recent study also described the activation of conserved ORs by short chain CAs using orthologs in *Aedes aegypti* and *Aedes albopictus* (Huff & Pitts, 2020). Lactic acid is a volatile CA that is emitted at a higher concentration in human skin as compared with non-human animals and is a powerful synergistic attractant of female mosquitoes (Acree *et al*. 1968; Dekker *et al*. 2002; Eiras & Jepson 1991; McMeniman *et al*. 2014; Siju *et al*. 2010; Steib *et al*. 2001). As such, lactic acid may convey species-level information that influences the innate preference for human blood meals in *Aedes* mosquito species (Acree *et al*. 1968; Dekker *et al*. 2002; Eiras & Jepson 1991; McMeniman *et al*. 2014; Siju *et al*. 2010; Steib *et al*. 2001). Importantly the IR8a coreceptor is required for CA responses in *Ae. aegypti*, as *Ir8a* mutant animals display both reduced electrophysiological responses and diminished attraction to CAs, including lactic acid (Raji *et al*. 2019). A number of CAs, including straight chain and branched CAs, may also have a positive impact on host-seeking and oviposition site selection in female mosquitoes (Day 2005; Dekker *et al*. 2002; Ponnusamy 2008, 2010a; Navarro-Silva 2009; Sivakumar 2011). Recent studies have demonstrated that individual differences in human carboxylic acid signatures impact attractiveness to female mosquitoes (De Obaldia *et al*. 2022; Omolo *et al*. 2021). Interestingly, differential attractiveness of human hosts was retained even when the coreceptors *Ir8a, Ir25a*, and *Ir76* were each genetically ablated in *Ae. aegypti*, despite overall reductions in host seeking behavior (De Obaldia *et al*. 2022).

Although the influence of IRs on mosquito attraction has been established, the receptive properties across the large families of IRs encoded in mosquito genomes remains an open question. One study demonstrated that a receptor from *Anopheles gambiae, Ir75k* (*AgamIr75k*), responds robustly to octanoic and nonanoic acids (Table 1; Pitts *et al*. 2017). In addition, members of the *Ir75* subfamily are responsible for the detection of CAs in the model fly, *Drosophila melanogaster* (Table 1; Benton *et al*. 2009; Ai *et al*. 2010; Abuin *et al*. 2011; Silberling *et al*. 2011; Prieto-Godino *et al*. 2016, 2017) and in the turnip moth, *Agrotis segetum* (Hou *et al*. 2022). The conservation and ancient origin of IRs suggests that orthologs of these receptors function similarly in other insects (Croset *et al*. 2010; Rytz *et al*. 2013). Interestingly, their potential roles in modulating host-seeking behaviors is supported by the down regulation of *Ir75* transcripts following a blood meal (Taparia *et al*. 2017). In the present study, we investigated the functional activation of receptors from the IR75 subfamily that are encoded in the genomes of two prominent vector species, *Ae. aegypti* and *Ae. albopictus*, using a heterologous expression platform. The orthologous receptors, *AaegIr75k1* and *AalbIr75e*, were maximally activated by nonanoic acid and to a lesser degree by octanoic acid. In contrast, a closely related receptor, *AaegIr75k3*, displayed activation responses to both octanoic and nonanoic acids that were similar magnitude and sensitivity, mimicking the activation profile of *AgamIr75k* (Pitts *et al*. 2017). All three *Aedes* receptors produced half-maximal effective concentrations in the low micromolar range, consistent with the hypothesis that CAs are their cognate ligands. Our results establish a foundation for functional genetic studies of the IR75 subfamily of receptors and their effects on mosquito behavior.

**Table 1.** Functional responses of IR75 subfamily receptors in flies.

| Receptor | Functional Response | References |
| --- | --- | --- |
| DmelIR8a | coreceptor | Benton *et al.* 2009 |
| AaegIR8a | coreceptor | Raji *et al.* 2019 |
| DmelIR84a | phenylacetic | Benton *et al.* 2009; Grosjean *et al.* 2011; Silbering *et al.* 2011 |
| DmelIR64a | various | Ai *et al.* 2010, 2013; Silbering *et al.* 2011; Gao *et al.* 2015 |
| DmeIR31a | 2-oxopentanoic | Benton *et al.* 2009; Silbering *et al.* 2011 |
| DmelIR75a | acetic, propionic | Benton *et al.* 2009; Abuin *et al.* 2011; Silbering *et al.* 2011; Prieto-Godino *et al.* 2016, 2017 |
| DmelIR75b/c | butyric, propionic, 2-oxopentanoic | Benton *et al.* 2009; Abuin *et al.* 2011; Silbering *et al.* 2011; Prieto-Godino *et al.* 2017 |
| AgamIr75k | octanoic, nonanoic | Pitts *et al.* 2017 |
| AaegIR75k1 | nonanoic | this study |
| AaegIR75k3 | octanoic, nonanoic | this study |
| AalbIR75e | nonanoic | this study |

## Methods

### Gene Annotations & Phylogenetic Analysis

IR75 homologs were identified in the genomes of Culicidae species via tBLASTn or BLASTp searches against available genome assemblies on the National Center for Biotechnology Information (NCBI; www.ncbi.nlm.nih.gov) or VectorBase (www.vectorbase.org). Gene annotations were corrected based upon multiple amino acid alignments using Geneious Prime^2019^ software (Biomatters Limited, USA) and conservation of intron positions. Alignment parameters were set for the MUSCLE algorithm with UPGMB clustering and 1000 pseudoreplicates (Edgar 2004a, 2004b). Branches having >50% bootstrap support were retained.

### IR gene expression analysis

Publicly available antennal RNAseq reads (Pitts *et al*. 2011; Lombardo *et al*. 2017; Matthews *et al*. 2016; Leal *et al*. 2013) were downloaded from NCBI and mapped to transcript assemblies obtained from VectorBase (*Aedes aegypti* LVP_AGWG, *Aedes albopictus* Foshan FPA, *Anopheles gambiae* PEST, *Culex quinquefasciatus* Johannesburg) into which new IR annotations were incorporated. Reads were mapped using the Sailfish alignment-free quantification software, with k-mers of 21 bp and sequence-specific bias correction (Patro *et al*. 2014) and expressed in transcripts per kilobase per million reads (TPM).

### Gene cloning and sequencing

Coding regions for AalbIR8a, AalbIR75e, AaegIR8a, AaegIR75k1, and AaegIR75k3 were de novo synthesized by Twist Biosciences (San Francisco, CA, USA), cloned into the pENTR^TM^ vector, and subcloned into the *Xenopus laevis* expression destination vector pSP64t-RFA using the Gateway^®^ directional cloning system (Invitrogen Corp., Carlsbad, CA, USA). Plasmids were purified using GeneJET Plasmid Miniprep Kit (ThermoFisher Scientific, Waltham, MA, USA) and sequenced in both directions to confirm completeness of coding regions.

### Chemical reagents

Chemicals used for the deorphanization of AaegIR75k1, AaegIR75k3 and AalbIR75e were obtained from Acros Organics (Morris, NJ, USA), Alfa Aesar (Ward Hill, MA, USA), and ThermoFisher Scientific (Waltham, MA, USA) at the highest purity available (Supplemental Table 1). Odorants were diluted into 1M stocks in 100% DMSO. Compounds were grouped by chemical classes and mixed at equimolar concentrations of [10^−4^ M] in ND96 buffer (96mM NaCl, 2mM KCl, 5mM MgCl_2_, 0.8mM CaCl_2_, and 5mM HEPES, pH 7.6).

### Two-electrode voltage clamping

*cRNA transcripts were synthesized from XbaI-linearized pSP64t-RFA expression vectors* using the mMESSAGE mMACHINE ^®^ SP6 kit (Life Technologies). Stage V-VII *Xenopus laevis* oocytes were purchased from Xenopus1 (Dexter, MI, USA) and maintained in incubation medium (ND96 96 with 5% dialyzed horse serum, 50μg/mL tetracycline, 100μg/mL streptomycin, 100μg/mL penicillin, and 550μg/mL sodium pyruvate) at 18^°^C. Oocytes were injected with 27.6 nL of each cRNA using a Nanoliter 2010 injector (World Precision Instruments, Inc., Sarasota, FL, USA). Odorant-induced currents were measured using the two-microelectrode voltage-clamp technique (TEVC) using an OC-725C oocyte clamp (Warner Instruments, LLC, Hamden, CT, USA) at −80mV holding potential. Blends were perfused over oocytes in ND96 buffer for 8-15 seconds. Inward currents were allowed to return to baseline prior to subsequent stimulus. Data acquisition was carried out with the Digidata 1550 B digitizer and pCLAMP10 software (Molecular Devices, Sunnyvale, CA, USA). Tuning curves were determined for 13 unitary carboxylic acids [10^−4^M]. All data analyses were performed using GraphPad Prism 8 (GraphPad Software Inc., La Jolla, CA, USA). Establishment of concentration-response curves were generated by exposing oocytes to octanoic or nonanoic acid (10^−9^ M to 10^−4^ M). To measure the effect of the compounds on the oocytes, odorants were perfused for up to 30s or until peak amplitude was reached. Current was allowed to return to baseline between chemical compound administrations. Raw data from oocyte recordings is provided in Table S2.

## Results

### IR75 orthologs

A conserved subfamily of variant ionotropic glutamate receptors (IRs) is encoded in the genomes of the vector mosquito species *Ae. aegypti*, *Ae. albopictus*, *An. gambiae*, and *Culex quinquefasciatus*, as well as the vinegar fly, *Drosophila melanogaster* (Figure 1 and Supplemental Table 2). While many of these IRs have been previously described, we have refined their annotations based upon amino acid similarities and conservation of introns (Figure 2; Supplemental Table 2). An example of the high degree of conservation and apparent IR orthology is the receptor pair AaegIR75k1 (AAEL014089) and AalbIR75e (AALFPA056866) that are 83% identical at the amino acid level and share the same set of introns (Figure 2). An apparent paralog of AaegIR75k1, AaegIR75k3 (AAEL023538), is also evident within the *Ae. aegypti* genome. The two receptors are 87% identical (Figure 2). These two genes are also closely linked (Supplemental Table 2), and share conserved intron positions (Figure 2), suggesting a potential recent duplication event. Additionally, we identified conserved receptors in *An. gambiae*, AgamIR75k (AGAP 007498) that has been functionally characterized (Pitts *et al*., 2017), and *Cx. quinquefasciatus*, CquiIR75k whose function is unknown (Figure 2). We also observed a group of nine CquiIR75 receptors (Figure 1) that are clustered in the *Cx. quinquefasciatus* genome, suggesting a common ancestral lineage and recent gene expansion (Supplemental Table 2). Members of the IR75 subfamily are generally expressed in the antennae of vector mosquitoes, consistent with their roles in olfactory sensory neuron function (Figure 3). A recent study indicated that IRs are broadly co-expressed with other chemoreceptor classes, especially ORs, in olfactory receptor neurons in the antenna (Herre et al. 2022). However, the overlap between IR8a and ORco appears to be minimal and the potential co-expression of IR75 subfamily receptors with ORs in antennal neurons remains unclear (Herre *et al*. 2022).

**Figure 1.**
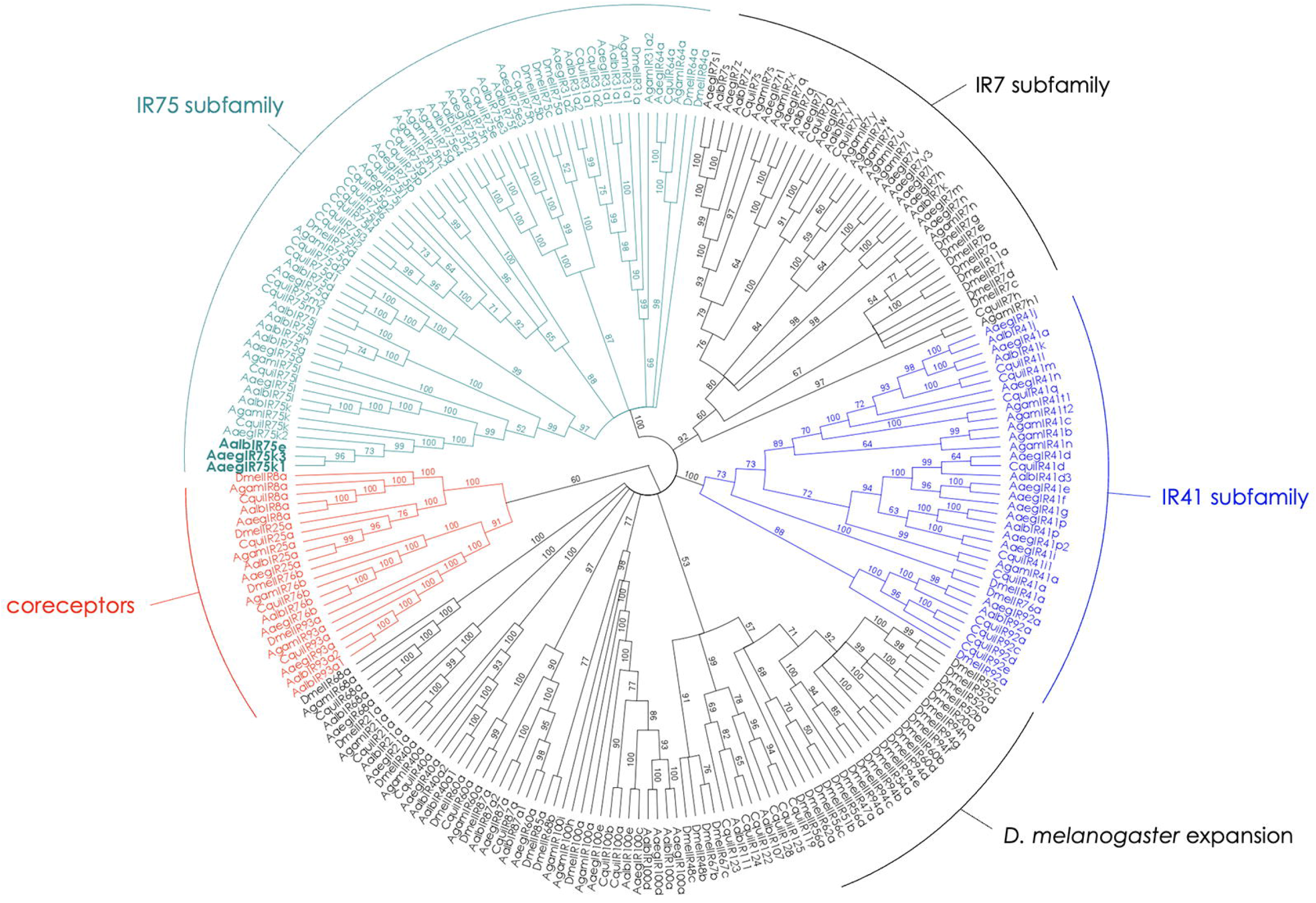
Variant Ionotropic Glutamate Receptors in Flies. Phylogenetic relationships among IR receptor proteins with bootstrap support for branches. IR75, IR7, and IR41 subfamilies are indicated, as well as an IR expansion in *D. melanogaster*. Coreceptors are highlighted in red.

**Figure 2.**
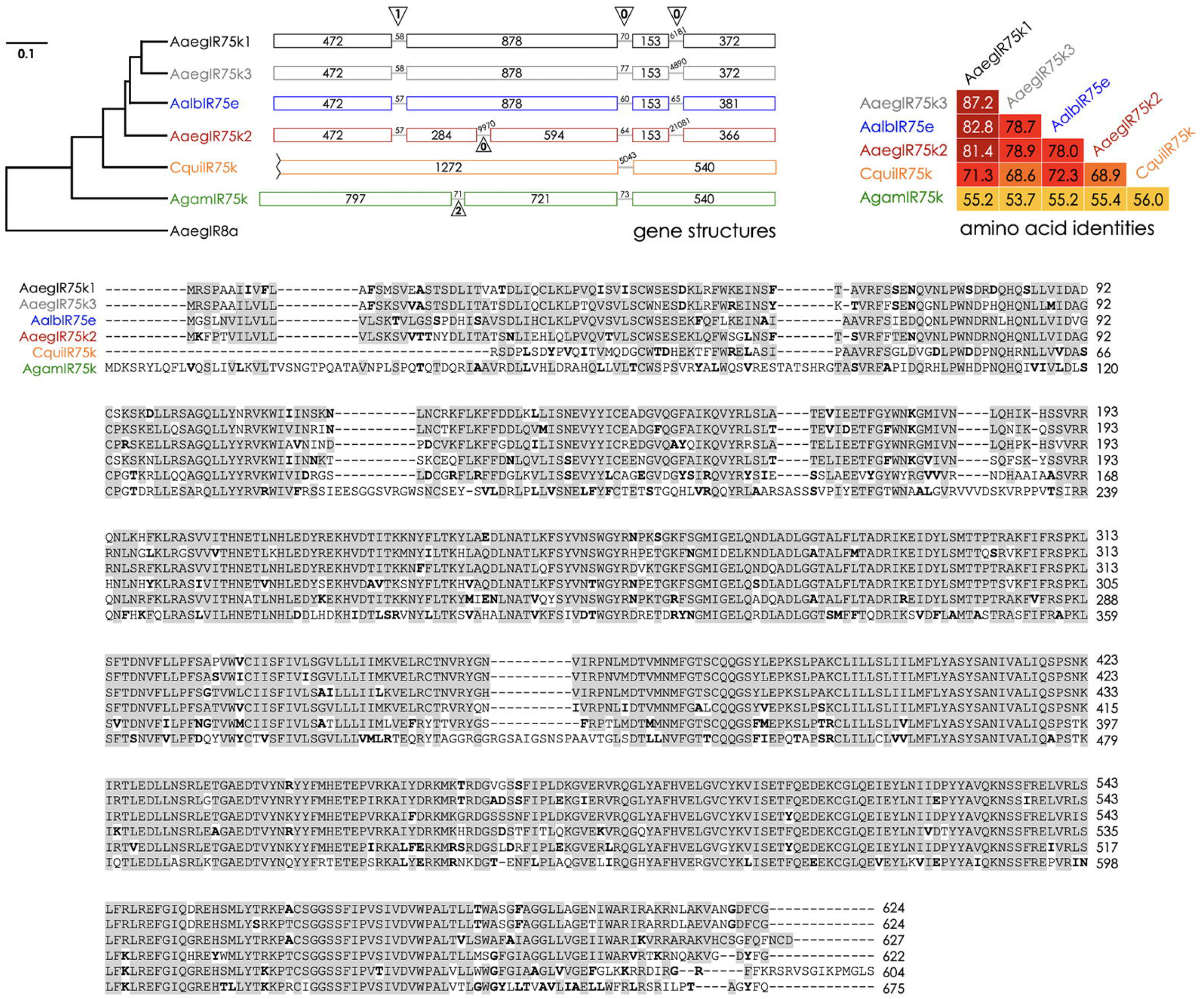
Highly Conserved IR75 Receptors in Vector Mosquitoes. *Top Panel:* Gene structures of IR75 receptors from *Ae. aegypti* (Aaeg), *Ae. albopictus* (Aalb), *Cx. quinquefasciatus* (Cqui), and *An. gambiae* (Agam). AaegIR8a is shown as an outgroup. Amino acid identities are provided for each receptor pair. *Bottom panel:* Amino acid alignment of IR75 receptors (single letter code). Gray boxes indicate identical amino acids while bold letters highlight conserved residues.

**Figure 3.**
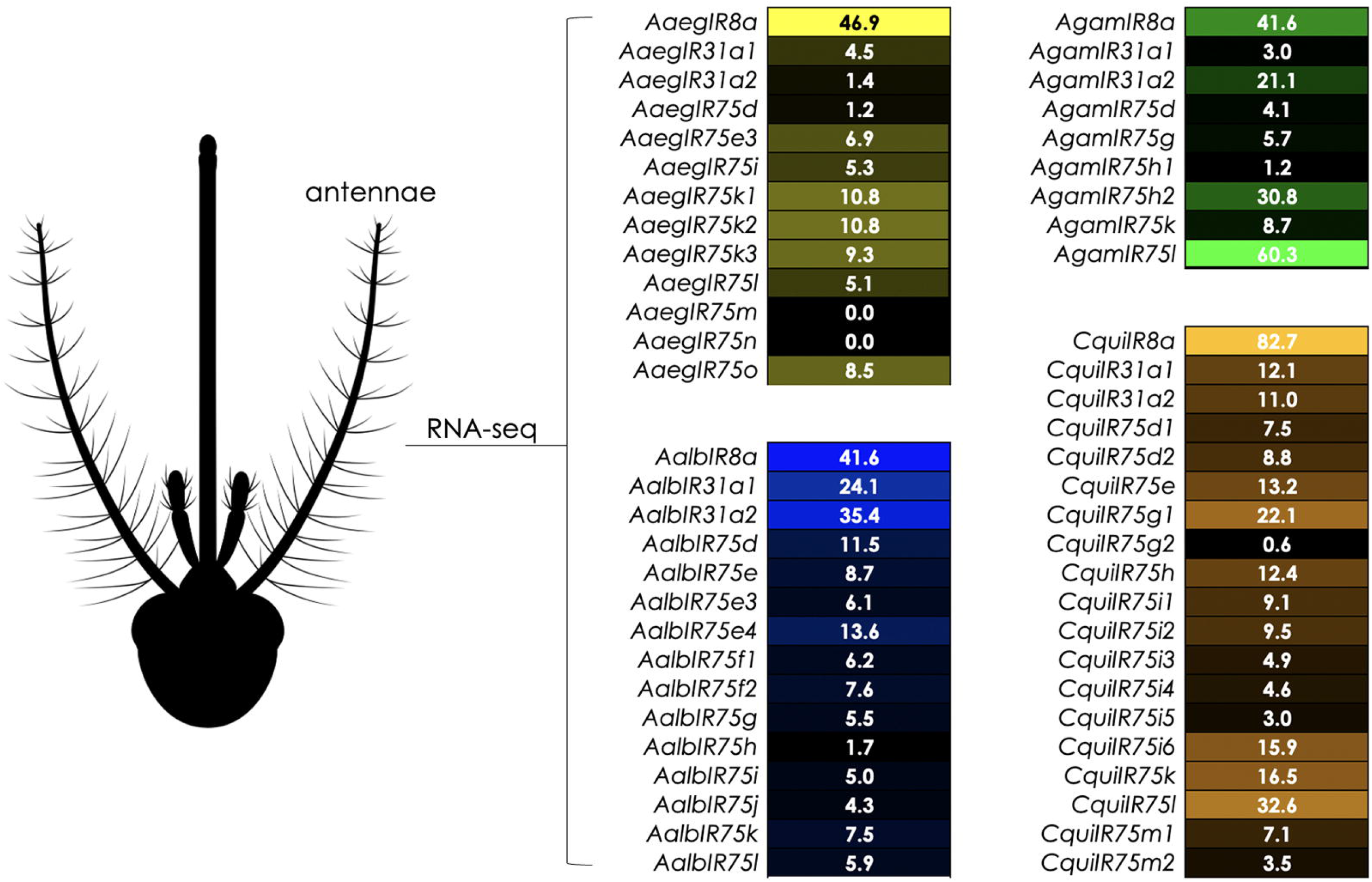
Expression of IR75 Transcripts in Female Antennae. Relative expression of IR75 subfamily members in Transcripts Per Million (TPM) in *Ae. aegypti* (yellow), *Ae. albopictus* (blue), *An. gambiae* (green) and *Cx. quinquefasciatus* (orange). IR8a coreceptor expression provided for comparison. Data derived from antennal RNAseq samples as described in: Pitts *et al*. 2011; Lombardo *et al*. 2017; Matthews *et al*. 2016; Leal *et al*. 2013.

### IR75 receptors are activated by carboxylic acids

Oocytes expressing receptor complexes consisting of an odor-tuning IR75 and the conspecific IR8a coreceptor were challenged with blends of compounds that were grouped by chemical structures, comprising at total of 73 individual compounds (Supplemental Table 1). AaegIR75k1, AaegIR75k3, and AalbIR75e all showed strong selective responsiveness to CAs when complexed with IR8a (Figure 4). Oocytes injected with single IR subunits did not respond to any of the tested compounds (data not shown). Receptor responses to CAs were approximately 10-fold greater than to any other odorant blend. The CAs that composed the blend were then tested individually at 10^−4^M. The AalbIR75e/AalbIR8a receptor pair responded maximally to nonanoic acid, with a magnitude that was three-fold higher than for octanoic acid (Figure 4). The AaegIR75k1/AaegIr8a receptor complex was also maximally activated by nonanoic acid, with responses four-fold greater than for octanoic acid (Figure 4). Interestingly, AaegIR75k3 elicited a dual selectivity to both octanoic and nonanoic acids (Figure 4). Responses to octanoic acid and nonanoic acid were approximately equal to one another and four-fold greater than the responses to any other CA. This AaegIR75k3 response profile is similar to AgamIR75k (Pitts *et al*. 2017).

**Figure 4.**
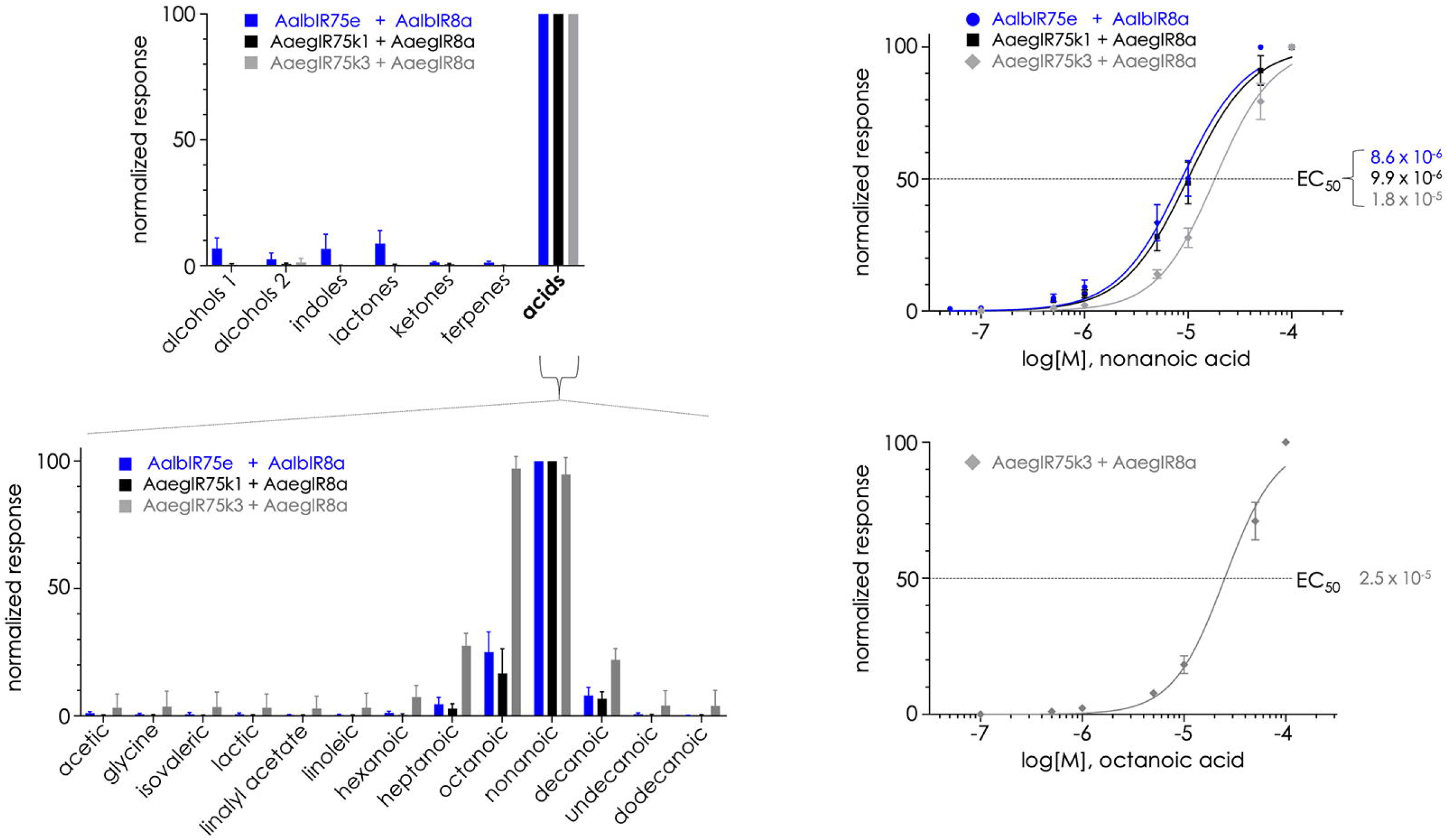
Odorant-Induced Electrophysiological Responses to Carboxylic Acids. Oocytes were clamped at −80mV and inward currents were normalized to the highest response. *Top left:* compound blends were arranged by chemical class at 10^−4^M and presented to oocytes. AalbIR75e, AaegIR75k1, and AaegIR75k3 responded maximally when exposed to the blend of carboxylic acids. *Bottom left*: individual compounds within the CA blend were perfused at 10^−4^M. Maximal activation of AalbIR75e (blue bars) and AaegIR75k1 (black bars) occurred with nonanoic acid. AaegIR75k3 (gray bars) responded with similar magnitudes to nonanoic and octanoic acid. *Top right:* Concentration response curves for nonanoic acid. The low micromolar range EC_50_ suggests that nonanoic acid is the cognate ligand of these receptors. *Bottom right*: Response curve for AaegIR75k3 and octanoic acid.

### Concentration responses of IR75 receptors

Responses of receptor complexes to concentrations of nonanoic acid ranging from 10 M to 10^−4^M (Figure 4) were measured. The resulting electrophysiological responses were fitted to sigmoid curves and half-maximal effective concentration values (EC_50_) were calculated for the CAs (Figure 4). All three receptors produced concentration dependent responses to CAs, consistent with ligand-gated ion channel openings at the membrane surface that are saturated at the highest concentrations tested (Figure 4). Nonanoic acid produced EC_50_ values of 8.64μM for AalbIR75e and 9.94μM for AaegIR75k1 (Figure 4). AaegIR75k3 responded to nonanoic acid with an EC_50_ value of 18.12μM, roughly 2-fold higher than the potency of nonanoic acid for AaegIR75k1. Because AaegIR75k3 was also activated by octanoic acid, we assayed concentration dependency and determined the EC_50_ value to be 25.16 μM, slightly higher than for nonanoic acid (Figure 4). Additionally, we observed that a heterospecific IR combination, AaegIR75k1/AalbIR8a, was also activated by nonanoic acid, although the EC_50_ value was approximately 2-fold higher than for the conspecific combination of AaegIR75k1/AaegIR8a (Supplemental Figure 1).

### Ionotropic Receptor Coreceptor Specificity

In an effort to demonstrate the sequence conservation and importance of *Ir8a* between mosquito species we performed experiments using heterospecific receptor partners from *Ae. aegypti* and *Ae. albopictus*. We utilized a combination of a tuning receptor from *Ae. aegypti* (*AaegIr75k1*) and the heterospecific coreceptor from *Ae. albopictus* (*AalbIr8a*) and determined the heterospecific receptor complex response profile to blends of chemicals as previously performed. The recorded responses show a strong activation to the CA blend and were similar to those of *AaegIr75k1* with the conspecific IR8 (Supplemental fig 1). The heterospecific receptor complex demonstrated a strong responsiveness to nonanoic acid (Supplemental fig 1). Additionally, the receptor complex showed a concentration dependent response profile when responding to nonanoic acid, with an EC_50_ value of 18.38 μM (Supplemental fig 1).

## Discussion

Functional studies of mosquito IRs are generally lacking, with the exception of AgamIR75k in *An. gambiae*, which responded best to octanoic and nonanoic acids (Pitts *et al*. 2017). We have discovered that the receptors AaegIR75k1, AaegIR75k3, and AalbIR75e are also activated by CAs, representing the first odor-tuning IRs to be characterized in *Aedes* species, to our knowledge. AaegIR75k1 and AalbIR75e were selective for nonanoic acid, although reduced responses to octanoic and decanoic acids were also evident (Figure 4). EC_50_ values for AaegIR75k1 and AalbIR75e were in the micromolar range for nonanoic acid, suggesting that this compound may act as a cognate ligand for these receptors in natural environments (Figure 4). AaegIR75k3 responded with nearly equal selectivity and sensitivity to both octanoic and nonanoic acids (Figure 4). Collectively, the demonstration that multiple receptors within the IR75 subfamily respond to CAs supports the hypothesis that additional IR75 receptors are likely to be tuned to the same compound class. Notably, despite overall high amino acid identities among the receptors we tested, functional conservation is still evident for the less conserved receptors. For example, AgamIR75k shares just over 50% identity with AaegIR75k1, AaegIR75k3, and AalbIR75e (Figure 2), yet all are activated by octanoic and nonanoic acids. The common sensitivities of IR75 receptors to octanoic and nonanoic acid across species leads us to speculate that short- to medium-chain CAs are of specific importance in the life histories of vector mosquitoes, potentially as host-seeking cues for females. We were also able to demonstrate that the response profile of a heterospecific receptor complex mimicked the conspecific receptor complex response (Supplemental Figure 1), implying that the IR8a coreceptor forms similar receptor complexes with tuning IRs in distinct species. One limitation of the heterologous oocyte cell system is that chemical compounds are delivered in an aqueous perfusion buffer, raising the possibility that solubility may affect receptor response amplitudes and therefore calculated EC_50_ values. We note that issues with volatility differences between compounds can also affect airborne stimulus delivery systems. Nonetheless, relative differences in efficacies and CRCs for each compound are still meaningful across species. Furthermore, identification of chemical receptors that respond to the same or similar cognate ligands is an important step toward determining environmental stimuli that are vital to the chemical ecology of vector mosquitoes.

Nonanoic and octanoic acids are key constituents of human sweat emanations, and are likely to impact behavioral outputs in female mosquitoes (Bernier *et al*. 2000; Cork & Park 1996; Dekker *et al*. 2002). Given their potential importance in female blood meal host seeking, expanding our analysis of the interactions between CAs and IR75 subfamily receptors will be important for enhancing our understanding of the chemical ecologies of vector species. Another aspect of these studies will be the regulation of IR75 receptors as a mechanism for modulating behavior. For example, one study found that three members of the IR75 subfamily in *Cx. quinquefasciatus*, *IR75e, IR75h*, and *IR75m2*, appeared to be downregulated following bloodfeeding (Taparia *et al*. 2017; Figure 3). Additionally, *AaegIR75k1* displayed reduced expression in the antennae of blood-fed females compared with sugar-fed or gravid individuals (Matthews *et al*. 2016), as well as in Dengue-infected vs. non-infected females (Tallon *et al*. 2020). In the latter study, Dengue infection led to reduced locomotion but increased sensitivity to human odors (Tallon *et al*. 2020). Interestingly, another study found differences in antennal olfactory gene expression across strains of *Ae. aegypti*, including members of the IR75 subfamily (Mitra *et al*. 2021). While the evidence is circumstantial, results like these may hint at differences in CA responses in adult females, depending on natural variations in populations or physiological states.

The genomic clustering of some IR75 receptors in mosquitoes also suggests possible adaptive radiations amongst members of this subfamily. Differential host selection between wildtype mosquitoes and *Ir75* mutants in a dual-choice olfactometer host selection paradigm could indicate that the Ir75 clade is implicated in the process of host-seeking and the degree of anthropophily displayed by some species (Takken & Verhulst 2013). Additional studies could investigate the role of various mosquito repellants, such as DEET, on the detection of CAs in *Ae. aegypti* and *Ae. albopictus*.

Previous studies in other insects, as well as those presented here, implicate IR75 subfamily receptors in the detection of CAs (Table 1 and references therein; Pitts *et al*. 2017; Hou *et al*. 2022). The conservation of receptor sequences and functionality across mosquito species, as well as in *D. melanogaster*, indicate the significance of environmental CA detection that is shared by dipteran flies and encoded in members of the IR75 subfamily. More specifically, the IR75 receptors that have been deorphanized in this study appear to be responsible, at least in part, for detecting nonanoic acid in mosquitoes (Figure 4). This finding does not rule out the possibility that other chemoreceptors encoded in the genomes of these species also contribute to the detection of nonanoic acid, as the broader tuning observed for AaegIR75k3 would indicate. By utilizing comparative functional studies on a larger scale, we expect that the accuracy of predicting chemosensory receptors will increase steadily. Moreover, as genetic modifications in mosquitoes become more accessible, correlating receptor responses in heterologous systems with *in vivo* phenotypes will provide valuable insights into neuronal physiology and natural behaviors such as host seeking and oviposition site selection.

---

**Supplemental Figure S1. Conspecific Versus Heterospecific IR Complexes**

*Left:* AaegIR75k1 in combination with coreceptors AaegIR8a (black) or AalbIR8a (green) responds with highest efficacy to nonanoic acid. *Right:* AaegIR75k responded with greater sensitivity to nonanoic acid in the conspecific configuration.

Supplemental Figure S2. IR75 receptors mediate responses to carboxylic acids in vector mosquitoes.

## Acknowledgements

We thank Shan Ju Shih (Baylor University) for technical assistance with oocyte maintenance and injections. This work was supported by the National Institutes of Allergy and Infectious Diseases at the National Institutes of Health [grant number R15 AI156684-01 to RJP].

## Author Contributions

RJP conceived the original study with recommendations from MD. RJP supervised experiments. GR and RMH conducted electrophysiological experiments. AJB and RJP mapped antennal RNAseq reads and produced transcript expression data. GR, RMH, JSC, AJB, MD and RJP analyzed data. GR, RMH and RJP drafted the manuscript and prepared figures. GR, RMH, JSC, AJB, MD and RJP revised the manuscript. All authors approved the final version of the manuscript.

## Notes

### Competing Interest Statement

The authors have declared no competing interest.

### Summary of Updates

Revised methods and references.

