## Supplementary figures and images for "Carboxylic Acids that Drive Mosquito Attraction to Humans Activate Ionotropic Receptors"

### Supplemental Figure S1

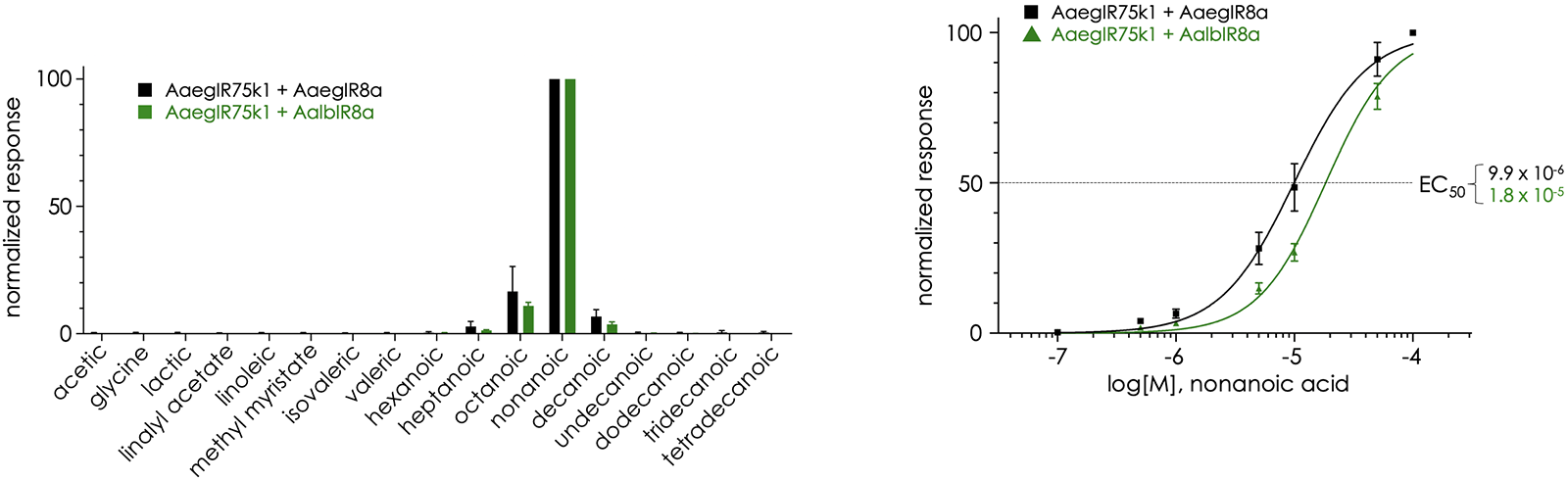

### Supplemental Figure S2

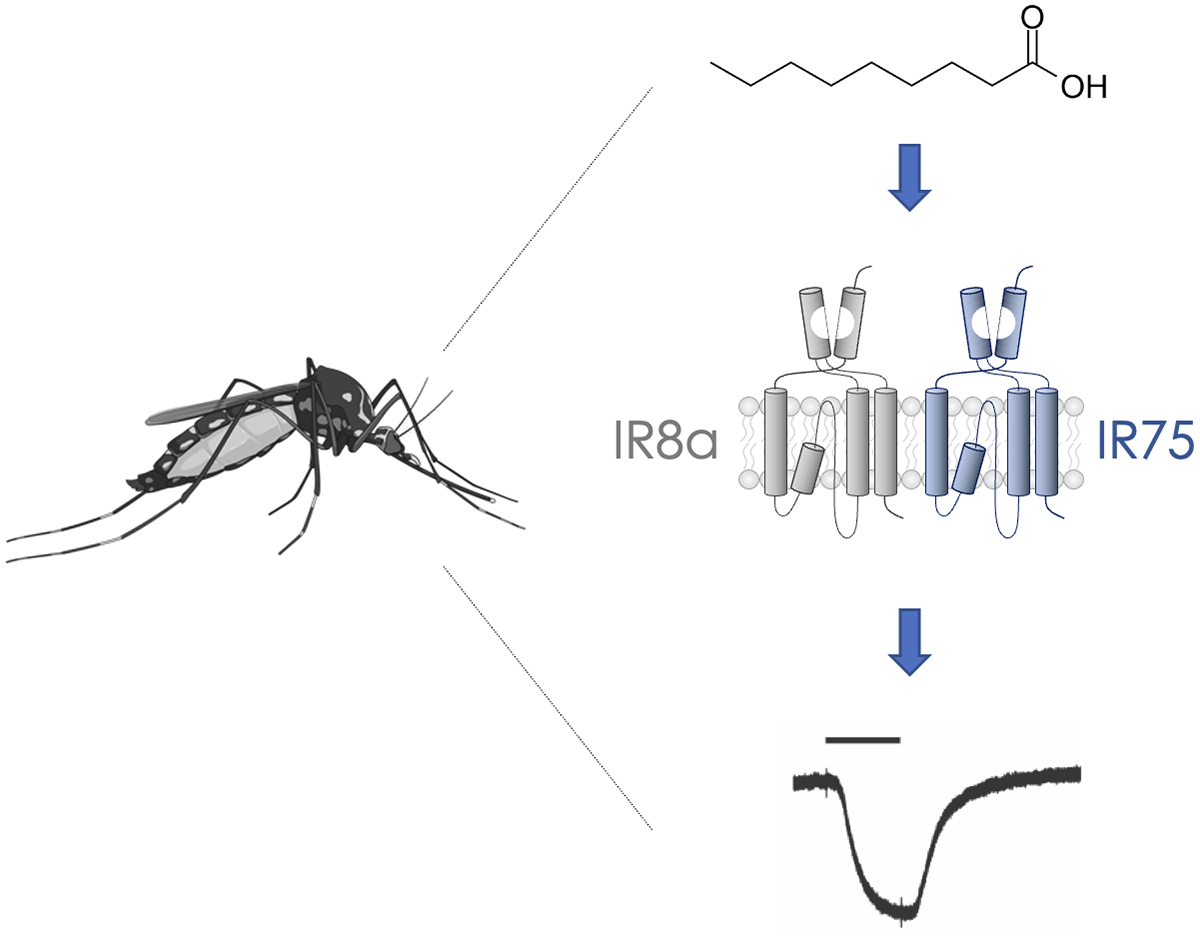
